# Predicting the current and future distribution of four original plants of Gentianae Macrophyllae Radix in China under climate change scenarios

**DOI:** 10.1101/2021.01.26.428226

**Authors:** Houkang Cao, Xiaohui Ma, Li Liu, Shaoyang Xi, Yanxiu Guo, Ling Jin

## Abstract

The wild resources of the four original plants (*Gentiana crasicaulis* Duthie ex Burk, *Gentiana daurica* Fisch, *Gentiana straminea* Maxim, and *Gentiana macrophylla* Pall) of Gentianae Macrophyllae Radix are becoming exhausted. Predicting the distribution under current and future climate scenarios is of significance for the sustainable utilization of resources and ecological protection. In this study, we constructed four species distribution models (SDMs) combining species distribution informations, 19 bioclimatic variables, and the maximum entropy (MaxEnt) model. The results showed that these 4 plants prefer a cool and humid climate. Under the future climate scenarios, the areas of the highly suitable habitats for *Gentiana crasicaulis* Duthie ex Burk and *Gentiana daurica* Fisch were likely to decrease, while *Gentiana straminea* Maxim was likely to expand, and *Gentiana macrophylla* Pall was less affected. In addition, the centroids of the highly suitable habitats for the four species shifted north or west. Most notably, most of the highly suitable habitats for the four species remained unchanged, which would be the preferred area for semi-artificial cultivation. The above information in this study would contribute to the development of reasonable strategies to reduce the impact of climate change on the four original plants.

## Introduction

The “Chinese Pharmacopoeia” stipulates that the original plants of Gentianae Macrophyllae Radix, called Qinjiao as a traditional Chinese medicine in China, are *Gentiana crasicaulis* Duthie ex Burk, *Gentiana daurica* Fisch, *Gentiana straminea* Maxim, and *Gentiana macrophylla* Pall [1]. There are more than 100 kinds of Chinese patent medicines containing Gentianae Macrophyllae Radix, which are used for the treatment of cholecystitis, hepatitis, psoriasis, rheumatoid arthritis and other diseases [2]. Extensive clinical application brings huge market demand, which leads to many wildly original plants being mined unreasonably and illegally. These four species were identified as national conserved wild plants in 1987, making artificial cultivation the only legal source of Gentianae Macrophyllae Radix, but the wild plants have not been effectively protected [3]. Although the government has been supporting the cultivation of original plants for more than 20 years, large-scale cultivation is still difficult. One of the most important reasons is the improper selection of suitable habitats. Therefore, predicting the current and future distribution of suitable habitats for original plants is an important measure for artificial cultivation.

Global warming is an indisputable fact. By the end of the 21st century, the average annual temperature will increase by 1.4-5.8 °C, which will further lead to changes in precipitation and other climate variables, thus leading to changes in the ecosystem [4]. More and more studies have shown that global warming can lead to the migration and diminishment of habitats, or even the extinction of some species due to lack of suitable habitats [5–8]. It is of great theoretical and practical significance for the effective protection of endangered species and biodiversity to predict the changes of suitable habitats for species under the future climate change scenarios [9–10]. Therefore, predicting the suitable habitats for the four original plants of Gentianae Macrophyllae Radix at present and in the future is of great value to formulating targeted measures as soon as possible.

Species distribution models (SDMs), which are jointly established by ecological niche model and future climate model, are widely used to predict the suitable habitats for species under the climate change scenarios [11–12]. This model first establishes the relationship between the distribution of current species distribution and climate variables, and then drives the model with the predicted scenarios of future climate change, so as to predict the response and distribution probability of species [13–14]. Among the many SDMs, the combination of maximum entropy (MaxEnt) model and Beijing climate center-climate system model (BCC-CSM) is very suitable for predicting the current and future distribution of Chinese species, and its prediction results are highly accurate, which has been proved in many studies on the distribution prediction of Chinese species [15]. Therefore, MaxEnt model and BCC-CSM-MR model were used in this study to predict the distribution of the four original plants of Gentianae Macrophyllae Radix.

In this research, we aimed to solve the following two problems: (1) To analyze the contribution of bioclimatic variables to SDMs and the corresponding optimal intervals. (2) To predict the area changes and centroid shifts of the highly suitable habitats under future climate scenarios, and find out the highly suitable habitats that would remain unchanged in the future.

## Methods

### Records of species distribution

In this study, we collected a total of 83 samples (17 *Gentiana crasicaulis* Duthie ex Burk, 26 *Gentiana daurica* Fisch, 19 *Gentiana straminea* Maxim and 21 *Gentiana macrophylla* Pall) from the wild, and all the samples had accurate longitude and latitude information. In addition, some distribution records with clear latitude and longitude were collected from the Chinese Virtual Herbarium database (CHV, http://www.cvh.ac.cn) and the Global Biodiversity Information Facility (GBIF, https://www.gbif.org/). All the longitude and latitude information of the samples (.csv format file) were imported into ArcMap 10.2 software. First, the “Projection and transform” function in “ArcToolbox” was used to convert the sample point information into “.shp” format. Then, the functions of “Buffer” and “Intersection” in “ArcToolbox” were used to screen out samples with a distance of less than 10km, which were eliminated to reduce the impact of sample bias [16]. Finally, the samples for subsequent prediction were determined, including 48 *Gentiana crasicaulis* Duthie ex Burk, 67 *Gentiana daurica* Fisch, 86 *Gentiana straminea* Maxim and 64 *Gentiana macrophylla* Pall (Fig 1).

**Fig 1.**
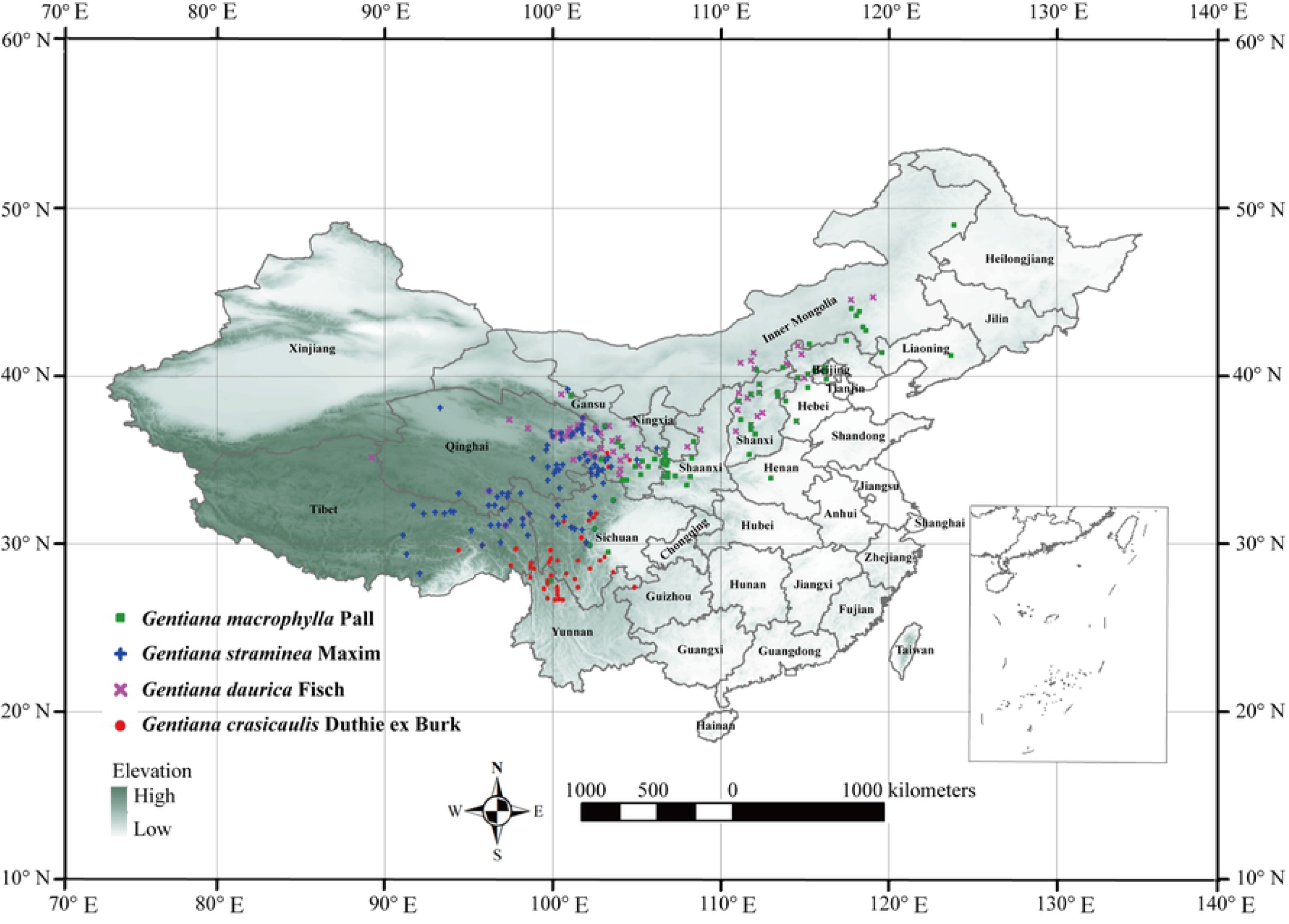
Distribution records of 4 species used for modeling in China

### Bioclimatic variable sources and data preprocessing

The current (1970-2000) and future climate data (2021-2100) including 19 bioclimatic variables were downloaded from the World Climate Database, with the resolution of 2.5 arcmin (https://www.worldclim.org/data/index.html, downloaded on July 23, 2020) [17]. In this study, the BCC-CSM-MR model (the upgraded version of BCC-CSM1.1) with strong simulation capability in China was selected as the future climate scenarios. The BCC-CSM-MR model published in the World Climate Research Program Coupled Model Intercomparison Project (CMIP6) contained four shared socioeconomic pathways (SSPs) SSP126, SSP245, SSP370 and SSP585, which replaced representative concentration pathways (RCPs) RCP2.6, RCP4.5, RCP6.0 and RCP8.5 in the CMIP5, respectively [18] (https://www.wcrp-climate.org/wgcm-cmip/wgcm-cmip6). Each scenario was divided into 4 future periods: the 2030s (2021-2070), the 2050s (2040-2060), the 2070s (2060-2080) and the 2090s (2080-2100). Firstly, according to the iteration operation results calculated in the MaxEnt 3.4.1 software based on the species latitude and longitude information and current climate variable data, the variables whose contribution was not 0 were selected. Secondly, the “Sampling” function of the “ArcToolbox” in the ArcMap software was performed to extract the values of the variables screened in the previous step. Thirdly, the Pearson correlations between the variables were conducted by SPSS 20.0 software, and the variables with a smaller contribution among the variables with a correlation coefficient greater than 0.8 were discarded to eliminate the negative impact of collinearity between variables on the model [19].

### Division and area of current and future habitats

The selected species distribution information and bioclimatic variables were imported into MaxEnt software for modeling operation, with the following parameter settings. 75% distribution data was used as a training set, and 25% was used as a test set [20]. The maximum number of backgrounds points was set to 10,000, and the maximum iterations was set to 500, with a 10-fold cross validation [21–23]. The area under the receiver operating characteristic (ROC) curve (AUC) was calculated to evaluate the accuracy of the model, and AUC values greater than 0.9 meant extremely high accurate [24]. In addition, the contributions of variables were determined by the jackknife analysis, and the optimal intervals of the variables were reflected by the response curves [25].

The logistic results of the ASCII document obtained by MaxEnt analysis are regarded as the probability of the species occurrence, and its value ranges from 0 to 1 [26]. In this study, the ASCII files were imported into ArcMap and converted into raster files [27]. The reclassification function was then performed to grade the habitats according to the recommendation of Intergovernmental Panel on Climate Change (IPCC) [28]: “not suitable habitat (*p* < 0.05)”, “lowly suitable habitat (0.05≤*p* <0.33)”, “moderately suitable habitat (0.33≤*p* <0.66)”, and “highly suitable habitat (*p* ≥0.66)”. By the property sheet of the classified file, the rate of the number of grids in each category to the total number of grids was calculated, which was equal to the area ratio. Then, the area of each habitat was calculated by the area ratio multiply by the total land area of China (960×10^4^ km^2^, http://www.gov.cn/guoqing/2017-07/28/content_5043915.htm).

### The centroids and unchanged area of the highly suitable habitats

The characteristics of the highly suitable habitat migration can be reflected by observing the shift of the centroids of the highly suitable habitats [29]. First, the raster layers of highly suitable habitats were extracted using the “Extract by attributes” function of “Spatial analysis tools” in “ArcToolbox”. Then, the centroids of the highly suitable habitats were determined using the “Zonal geometry” function in “Spatial analyst tools”.

Under future climate scenarios, areas that remained stable and unchanged in highly suitable habitats were analyzed with the help of “ArcToolbox”. First, all raster layers of highly suitable habitats were converted to surface layers (.shp files). Then, the “Clip” function in the “Analysis tools” was used to clip all the surface layers to obtain a surface layer, which was the stable and unchanged area.

## Results

### The model accuracy and the contributions and suitable intervals of bioclimatic variables in the current climate

From the 19 bioclimatic variables, 4, 6, 7 and 6 variables were screened respectively for the modeling of 4 species (Tab 1). The contributions of variables to each species calculated by jackknife analysis were as follows (Tab 1). The bioclimatic variables that contributed the most to distribution model of *Gentiana crasicaulis* Duthie ex Burk were isothermality (Bio3, 37.3%) and annual precipitation (Bio12, 31.5%). Precipitation of the driest month (Bio14, 31.5%) and precipitation of the warmest quarter (Bio18, 27.6%) made the greatest contributions to the distribution model of *Gentiana daurica* Fisch. The top two variables contributing to the distribution model of *Gentiana straminea* Maxim were mean temperature of the warmest quarter (Bio10, 44.6%) and precipitation of the warmest quarter (Bio18, 23.2%). Mean temperature of the driest quarter (Bio9, 24.8%) and precipitation of the warmest quarter (Bio18, 18.7%) contributed the most to distribution model of *Gentiana macrophylla* Pall.

**Tab 1.**
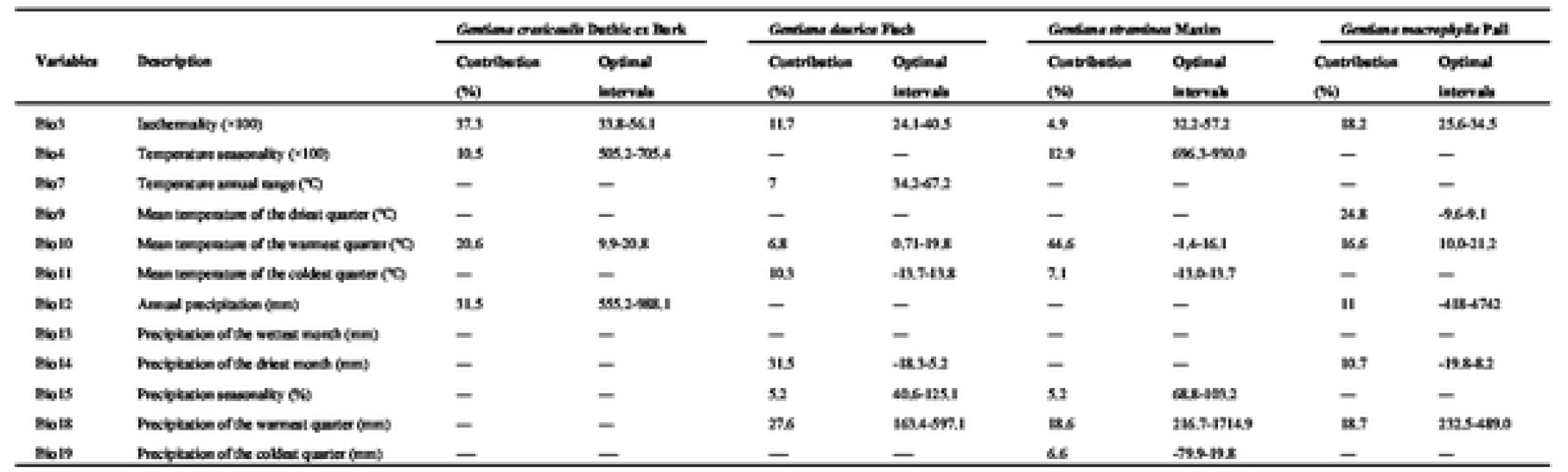
Percentage contributions and optimal intervals of the bioclimatic variables included in the Maxent models of 4 species

The relationship between the probability of species existence and the variables can be described by the response curves, which can also describe the optimal intervals for bioclimatic variables [30]. In this study, two bioclimatic variables isothermality (Bio3) and mean temperature of the warmest quarter (Bio10) common to the four species were observed. The optimal intervals of isothermality (Bio3) and mean temperature of the warmest quarter (Bio10) corresponding to *Gentiana crasicaulis* Duthie ex Burk, *Gentiana daurica* Fisch, *Gentiana straminea* Maxim and *Gentiana macrophylla* Pall were 9.9-20.8 °C, 0.71-19.8 °C, −1.4-16.1 °C and 10.0-21.2 °C, respectively (Tab 1).

### Prediction of the distribution of four species under current climate

At present, the highly suitable habitats for *Gentiana crasicaulis* Duthie ex Burk were mainly distributed in western Sichuan, northwest Yunnan and a small part of eastern Tibet, with a total area of 9.52×10^4^ km^2^, accounting for 0.99% of the total land area of China. The areas of moderately suitable habitats and lowly suitable habitats were 42.48×10^4^ km^2^ and 79.80×10^4^ km^2^, respectively (Fig 2).

**Fig 2.**
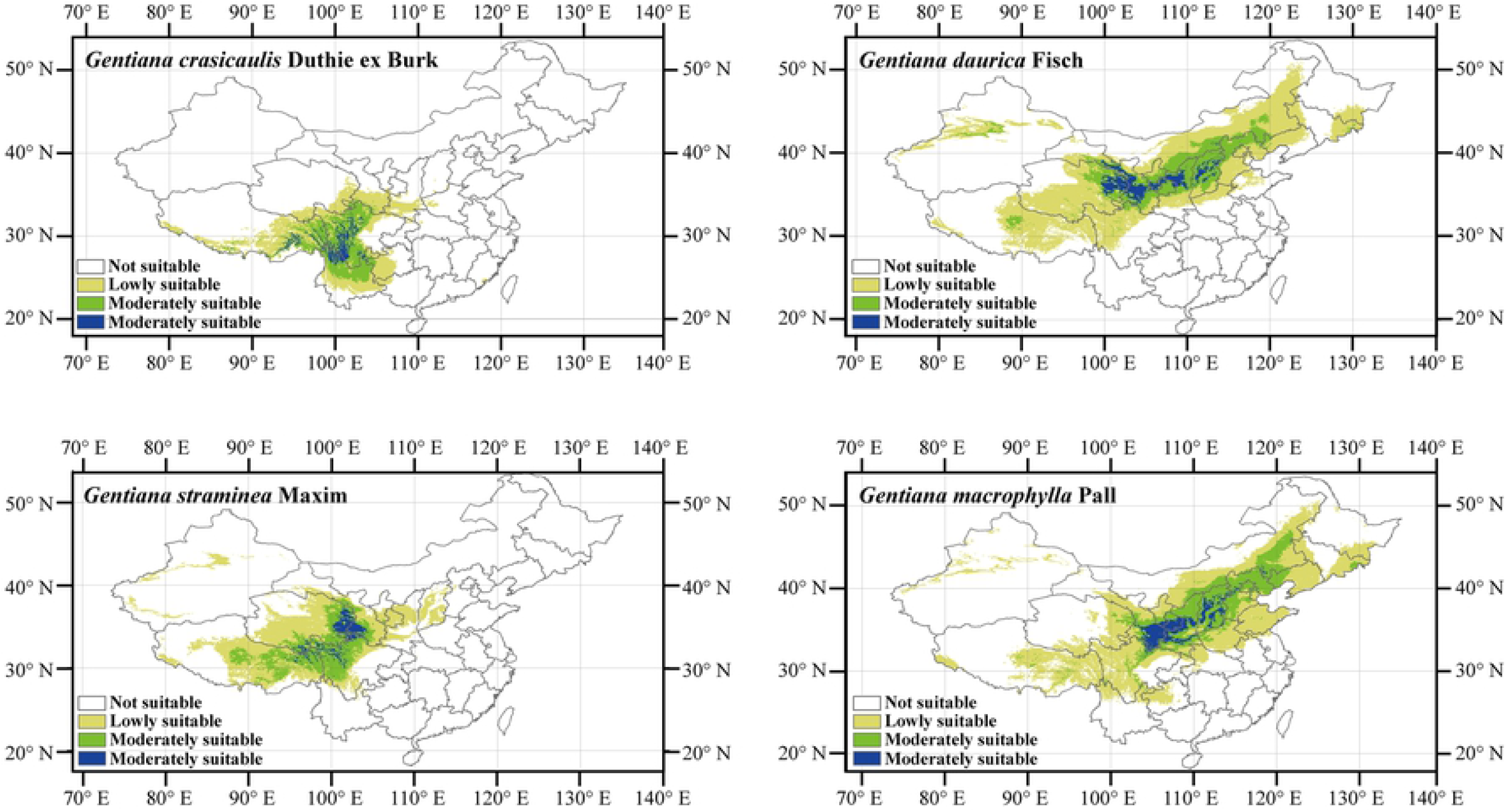
Prediction of the distribution of four species under current climate in China

The currently highly suitable habitats for *Gentiana daurica* Fisch existed in eastern Qinghai, southern Gansu, northwestern Shaanxi and central Shanxi, with a total area of 18.10×10^4^ km^2^, accounting for 1.89% of the total land area of China. The areas of moderately suitable habitatS and lowly suitable habitatS were 70.24×10^4^ km^2^ and 231.74×10^4^ km^2^, respectively (Fig 2).

Under the current climate, most of the highly suitable habitats for *Gentiana straminea* Maxim occurred in the eastern Qinghai, southern Gansu, and the sporadic part of northwest Sichuan and northeast Tibet, with a total area of 11.13×10^4^ km^2^, accounting for 1.16% of the total land area of China. The areas of moderately suitable habitats and lowly suitable habitats were 52.71×10^4^ km^2^ and 118.62×10^4^ km^2^, respectively (Fig 2).

The highly suitable habitats for *Gentiana macrophylla* Pall in the current were mainly distributed in southern Gansu, southern Ningxia, Shaanxi and central Shaanxi, with a total area of 19.07×10^4^ km^2^, accounting for 1.99% of the total land area of China. The areas of moderately suitable habitats and lowly suitable habitats were 76.92×10^4^ km^2^ and 223.51×10^4^ km^2^, respectively (Fig 2).

### Changes in the areas and centroids of the highly suitable habitats for four species and unchanged areas under future climate scenarios

#### *Gentiana crasicaulis* Duthie ex Burk

In terms of the degree of area change rates, the degree of change in both the highly suitable habitats and the moderately suitable habitats was relatively obvious under the future climate scenarios, while the degree of change in the lowly suitable habitats was slightly lower than that of the moderately suitable habitats, with the maximum increase or decrease of about 10%. Under the SSP245-2050s and SSP245-2080s scenarios, the area of highly suitable habitats increased by nearly 10%, but in different periods of SSP370 and SSP585, the area of highly suitable habitats showed a downward trend, with the maximum decline of nearly 10% (Fig 3A).

**Fig 3.**
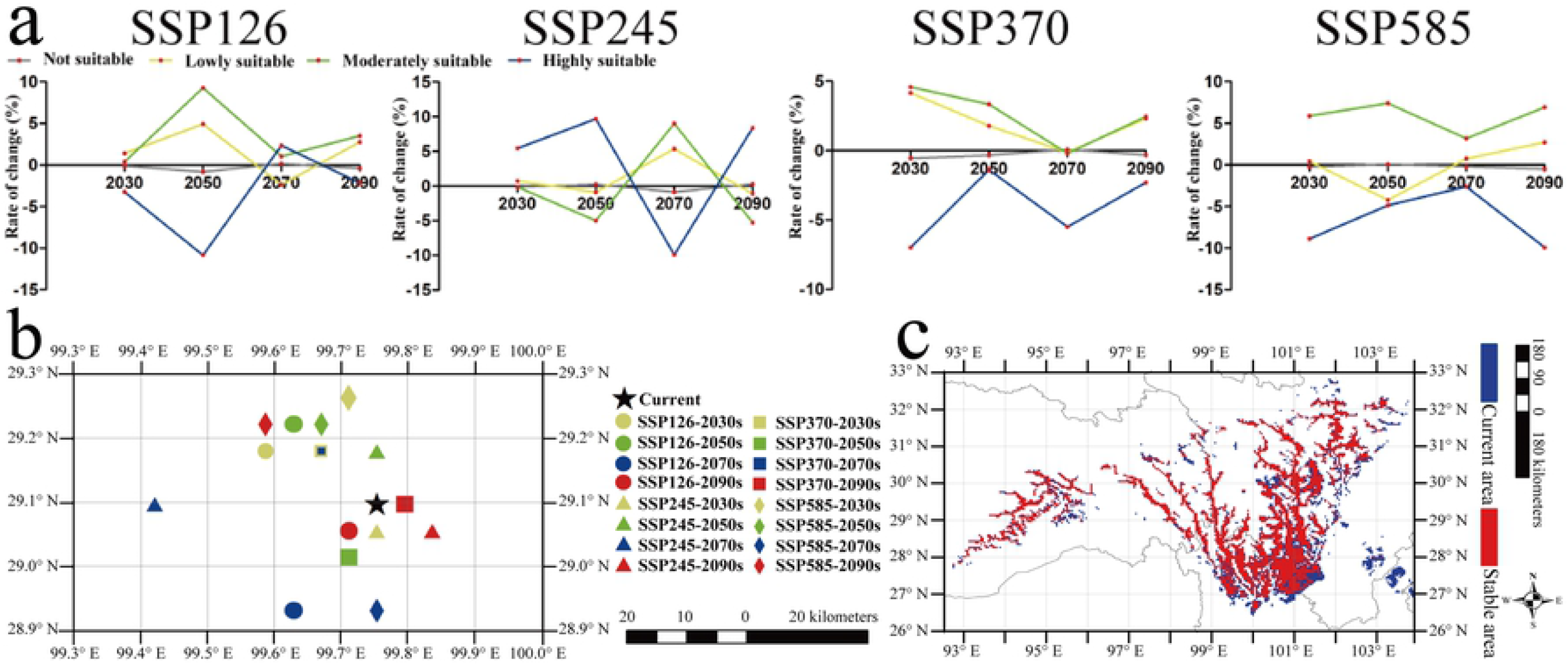
Changes in the areas and centroids of the highly suitable habitats for *Gentiana crasicaulis* Duthie ex Burk and unchanged areas under future climate scenarios

Compared with the current location of the centroid of the highly suitable habitats, the centroid shift directions in the future climate scenarios were mainly northwest and southwest, only the centroids under SSP245-2090s and SSP370-2090s scenarios shifted to the east, with the maximum shift distance of 37.07 km (SSP245-2070s) (Fig 3B). Under any future climate scenario, the area where the highly suitable habitats remained unchanged was 6.64 × 10^4^ km^2^, accounting for 69.69% of the current distribution area (Fig 3C). The stable habitat was mainly distributed in southwestern Sichuan.

#### *Gentiana daurica* Fisch

In terms of change trends, the moderately suitable habitats and the lowly suitable habitats were similar, and the highly suitable habitat and the not suitable habitats were similar, while the two change trends were roughly opposite. The area of highly suitable habitats showed a general trend of decline under most scenarios, and only showed a small trend of increase under SSP245-2050s scenario. Most of the area change rates were close to 5%, and the highest was about 10% (Fig 4A).

**Fig 4.**
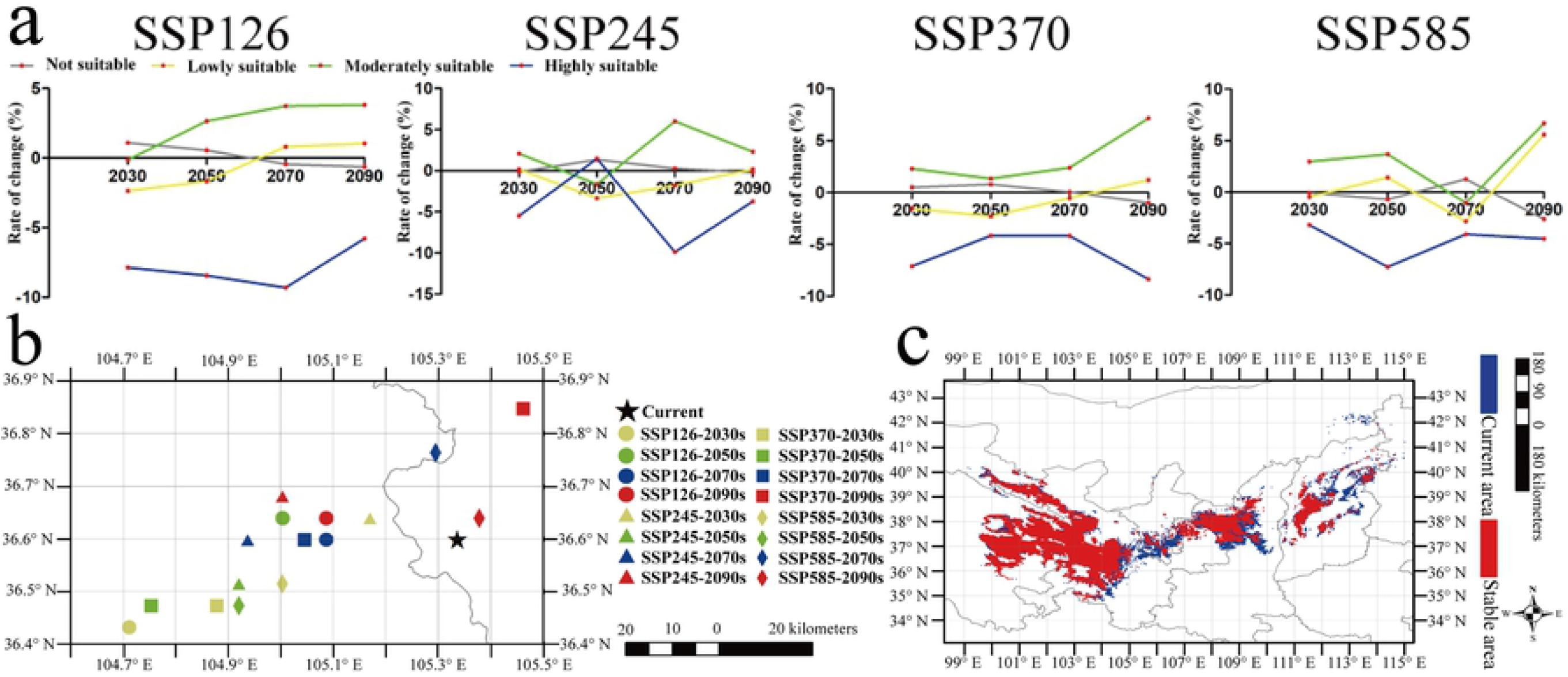
Changes in the areas and centroids of the highly suitable habitats for *Gentiana crasicaulis* Duthie ex Burk and unchanged areas under future climate scenarios

All centroids were distributed from northeast to southwest, and the future centroids shifted to the west or north of the current centroid, with a maximum shift distance of 73.27km (SSP126-2030s) (Fig 4B). The area of high suitable habitats that remained stable in the future was 12.83×10^4^ km^2^, accounting for 70.85% of the current distribution area (Fig 4C). The highly suitable habitats that remained unchanged occurred in the south of Gansu and the northwest of Shaanxi.

#### *Gentiana straminea* Maxim

The area changes of highly suitable habitats and moderately suitable habitats were obviously opposite, and the degree of increase or decrease could be as high as nearly 10%. On the whole, the highly suitable habitats showed an increasing trend with the highest that exceed 10% under most future climate scenarios, and it was slightly reduced only under the scenarios of SSP245-2070s and SSP370-2090s (Fig 5A).

**Fig 5.**
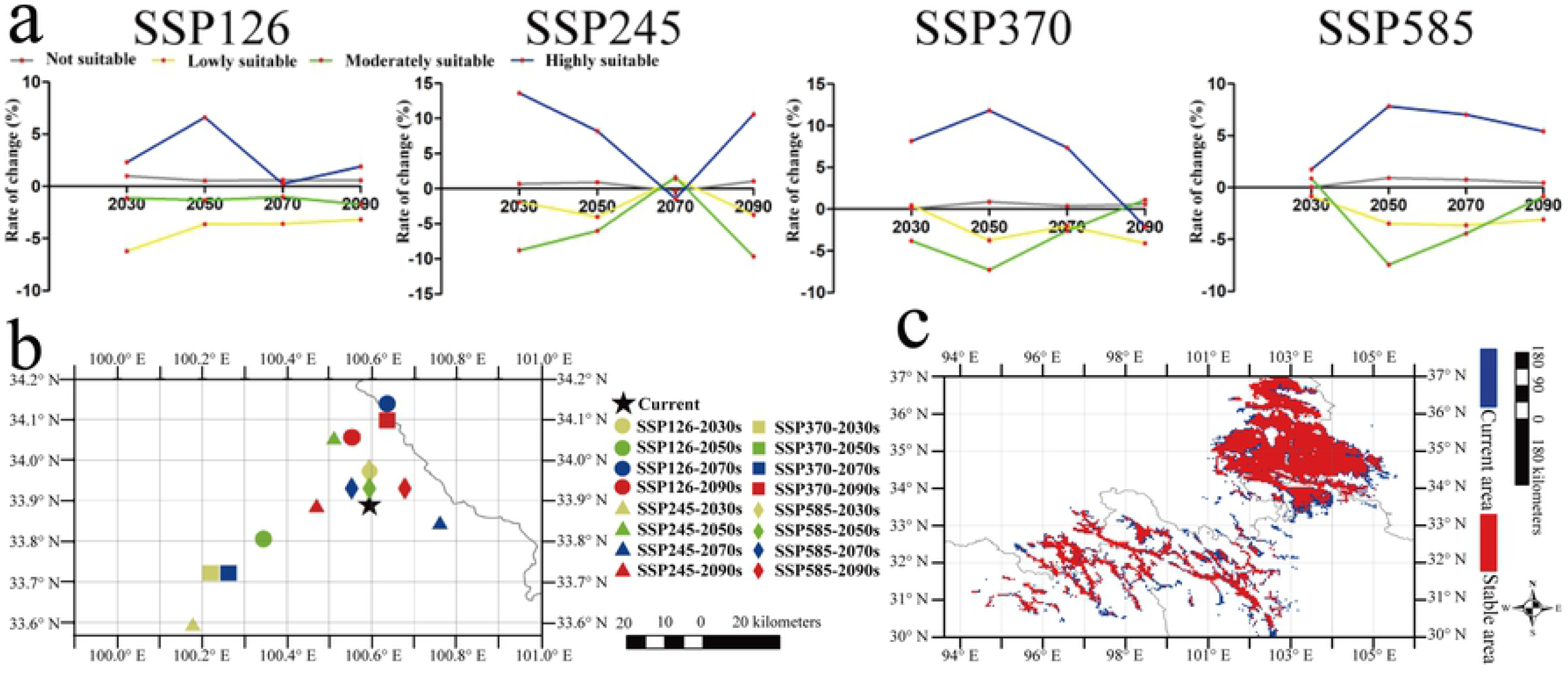
Changes in the areas and centroids of the highly suitable habitats for *Gentiana straminea* Maxim and unchanged areas under future climate scenarios

All centroids were roughly located from northeast to southwest. Compared with the current position of the centroid, most centroids of highly suitable habitats shifted to the north or southwest under the future scenarios, and the maximum shift distance was 60.51 km (SSP245-2030s) (Fig 5B). The area where the highly suitable habitat remained unchanged under the future climate scenarios was 8.31×10^4^ km^2^, accounting for 74.64% of the current area (Fig 5C). The highly suitable habitats that remained unchanged were mainly distributed in eastern Qinghai and southern Gansu.

#### *Gentiana macrophylla* Pall

Under the future scenarios, the change trend of highly suitable habitats for *Gentiana macrophylla* Pall was similar to that of moderately suitable habitats, but the change range of both was small, and the change rates of area were mostly within 5%. In addition, the possibility of expansion or contraction of the highly suitable habitats was similar, with greater volatility. Although the area was increased under the SSP585 scenarios, it did not exceed 4%. In general, the area of highly suitable habitats changed less under the future climate scenarios (Fig 6A).

**Fig 6.**
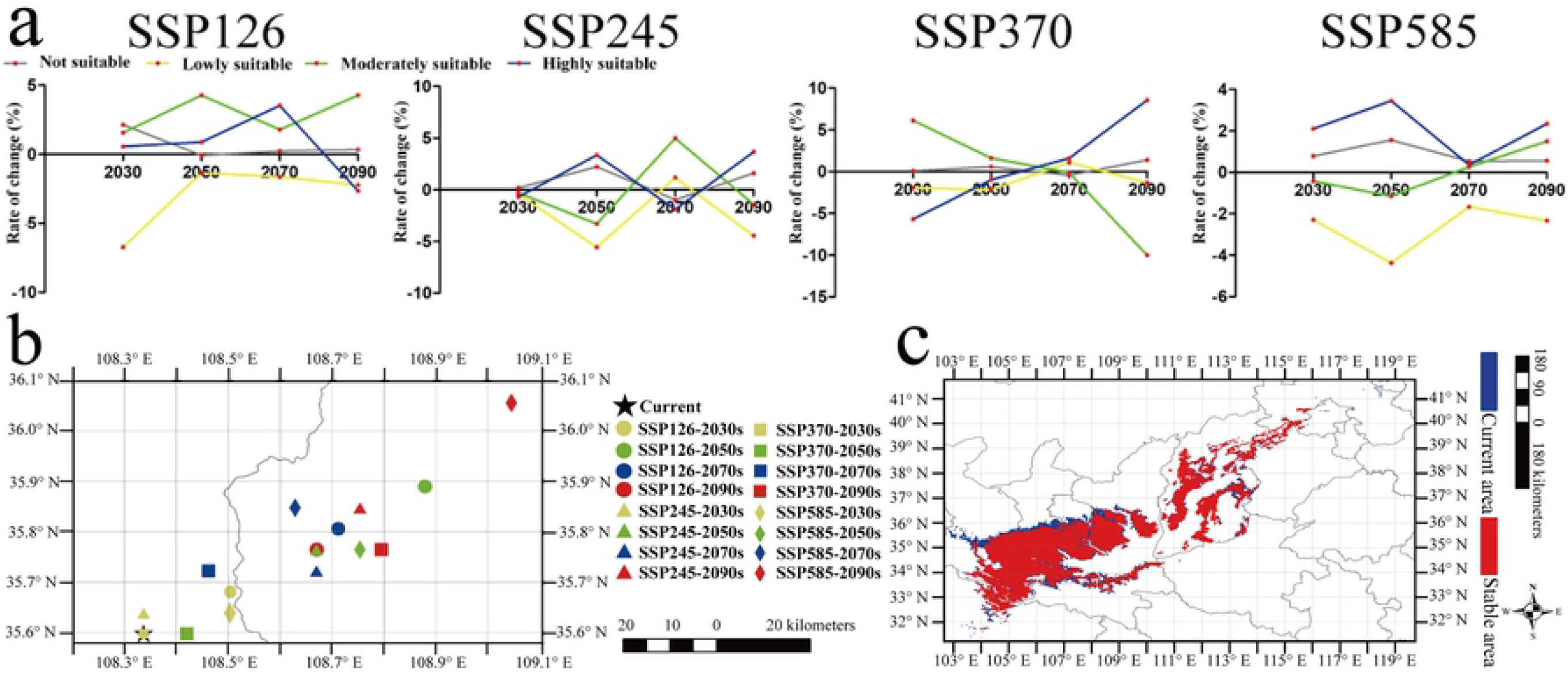
Changes in the areas and centroids of the highly suitable habitats for *Gentiana macrophylla* Pall and unchanged areas under future climate scenarios

All the centroids were distributed from northeast to southwest. Under the future climate scenarios, the current centroid mainly shifted to the northeast, and the maximum shift distance was 100.71 km (SSP585-2090s) (Fig 6B). The area of highly suitable habitats that remained stable under the future scenarios was 15.52×10^4^ km^2^, accounting for 81.37% of the current area (Fig 6C). The stable habitat was mainly distributed in southern Gansu.

## Discussion

The development, formation and distribution of plants are closely integrated with climate factors, especially temperature and precipitation [31]. First, we found that the variables contributing to the models of the four species were isothermality (Bio3) and mean temperature of the warmest quarter (Bio10). The optimal intervals of isothermality (Bio3) corresponding to *Gentiana crasicaulis* Duthie ex Burk, *Gentiana daurica* Fisch, *Gentiana straminea* Maxim and *Gentiana macrophylla* Pall were 33.8-56.1, 24.1-40.5, 32.2-57.2 and 25.6-34.5, respectively, which were at a medium to high level, which were conducive to the accumulation of chemical components to a certain extent [32]. Especially for *Gentiana crasicaulis* Duthie ex Burk, the contribution of isothermality (Bio3) was the highest compared with other variables, reaching 37.3%. In addition, the optimal intervals of mean temperature of the mean temperature of the warmest quarter (Bio10) were 9.9-20.8 °C, 0.71-19.8 °C, −1.4-16.1 °C and 10.0-21.2 °C, respectively. It was observed that the highest average temperature in the hottest season was only about 20 T, which was roughly at the medium temperature level. This was consistent with the characteristics of the four original plants distributed in low temperature areas in central and western China [33]. Especially for *Gentiana straminea* Maxim, the contribution of mean temperature of the warmest quarter was the highest compared with other variables, reaching 44.6%. This result was consistent with its location in the westernmost part of China among the 4 species, where the temperature was lower due to the high altitude [34]. In addition, the mean temperature of the mean temperature of the warmest quarter (Bio10) contributed the most to *Gentiana macrophylla* Pall (24.8%). It could be seen that moderate or low temperature had a great influence on the growth distribution of the four species, which was consistent with the seed germination experiments showing that the suitable temperature of the four species was below 25 °C [35–38]. As for the influence of precipitation, the contribution of annual precipitation (Bio12) to *Gentiana crasicaulis* Duthie ex Burk (31.5%) was slightly lower than isothermality. Precipitation of the driest month (Bio17) contributed the most to *Gentiana daurica* Fisch (31.5%). Precipitation of the warmest quarter (Bio18) contributed the most to *Gentiana straminea* Maxim and *Gentiana macrophylla* Pall (18.6% and 18.7%). Altogether, the precipitation had a greater impact on *Gentiana crasicaulis* Duthie ex Burk and *Gentiana daurica* Fisch, but a smaller impact on *Gentiana straminea* Maxim and *Gentiana macrophylla* Pall. According to the precipitation and geographical distribution of the highly suitable habitats of the four species, they are roughly located in the subhumid region of China, which conformed to the characteristics of their preference for wet climate [39–40]. To sum up, the habitats of the four species were characterized by dampness and coolness.

The core objective of this study was to predict the changes of suitable habitats (especially highly suitable habitats) for the four species under the future climate scenarios and develop corresponding strategies based on the changes. From the change rates of highly suitable habitat area, the change trends of the four species had certain differences. The distribution area of *Gentiana crasicaulis* Duthie ex Burk and *Gentiana daurica* Fisch generally showed a downward trend, especially under SSP370 and SSP585 scenarios. This result indicated that under higher temperature conditions, these two species may face the risk of shrinkage of highly suitable habitats, which was consistent with the challenges faced by many other plants [41]. However, climate warming does not mean that all species are facing a decline in suitable habitats, and there is also a situation of expansion of suitable habitats [42]. In the model of *Gentiana straminea* Maxim, the distribution of highly suitable habitats showed varying degrees of expansion at almost any period under any scenario. This result made us a little relieved about the future situation of *Gentiana straminea* Maxim. The change of *Gentiana macrophylla* Pall distribution area had obvious volatility, and the possibility of shrinkage and expansion was similar, but the magnitude of the change was small, which meant that the climate had little influence on the total area of the highly suitable habitats. In addition to the area changes, a feature of distribution changes, the influence of climate on the migration of suitable habitats for species is also important [43]. In this study, the future centroids of the four species generally displayed the characteristic of shifting north or west compared to the current position of the centroids, and the characteristic was the same as that of many species distributed in central and western China under the future scenarios [44–45]. The centroids of the four species had different shift distances in different periods under different scenarios, and the maximum centroid shift distances of each species ranged from 30 km to 100 km, which may be caused by the complex terrain and the suitable conditions of the species.

Although the highly suitable habitats of the species may change in area or centroid under future scenarios, some habitats remain unchanged [46]. In this study, we found that the area and location of the 70%-81% highly suitable habitats remained unchanged, and they were the habitats we focused on. Although the original plants of Gentianae Macrophyllae Radix had been planted in China for more than 20 years, it is difficult to promote artificial cultivation due to various reasons, one of which is the low germination rate and poor quality caused by improper selection of suitable habitat [47]. In order to solve this problem, semi-artificial cultivation is a promising approach by artificial seeding in the highly suitable wild habitats for species [48–49]. Based on the habitats for the original plants that remained stable and unchanged under future climate scenarios, we recommend artificial cultivation in the following habitats. Southwest Sichuan was suitable for *Gentiana crasicaulis* Duthie ex Burk, south Gansu and northwest Shaanxi were suitable for *Gentiana daurica* Fisch, east Qinghai and south Gansu were suitable for *Gentiana straminea* Maxim, and south Gansu was suitable for *Gentiana macrophylla* Pall. Therefore, this study suggested that semi-artificial cultivation of the four original plants in the highly suitable habitats was a reasonable measure to avoid the planting failure caused by future climate change. In addition, this measure could not only provide a legal source for Gentianae Macrophyllae Radix, but also help protect the ecological environment.

By establishing the SDMs, we found that the areas of highly suitable habitats for *Gentiana crasicaulis* Duthie ex Burk and *Gentiana daurica* Fisch were likely to decrease under future climate scenarios, while *Gentiana straminea* Maxim was likely to expand, and the distribution area of *Gentiana macrophylla* Pall changed less. In addition, most of the centroids of the highly suitable habitats for the four species shifted north or west. However, most of the highly suitable habitats for the 4 species remained unchanged under any future climate scenario, which would become the priority habitats for semi-artificial cultivation.

## Acknowledgments

The authors thank all researchers who contributed to this work by reporting distribution records of 4 species. This work was supported by the National Natural Science Foundation of China (81860673), Gansu Science and Technology Fundamental Innovation Project (1606RJIA323), and National Survey of Traditional Chinese Medicine Resources (Caishe [2018] No. 43).

## Author contributions

Data curation: Houkang Cao, Xiaohui Ma.

Formal analysis: Houkang Cao.

Funding acquisition: Ling Jin.

Investigation: Xiaohui Ma, Li Liu, Shaoyang Xi, Yanxiu Guo.

Methodology: Houkang Cao.

Project administration: Ling Jin.

Writing – original draft: Houkang Cao, Xiaohui Ma.

Writing – review & editing: Houkang Cao.

